# Thin-film freeze-drying of a Norovirus vaccine candidate

**DOI:** 10.1101/2021.06.08.447451

**Authors:** Haiyue Xu, Tuhin Bhowmik, Kevin Gong, Thu Ngoc Anh Huynh, Robert O. Williams, Zhengrong Cui

## Abstract

A bivalent Norovirus vaccine candidate has been developed that contains Norovirus strain GI.1 Norwalk-virus like particles (VLP) and strain GII.4 Consensus VLP adsorbed onto aluminum (oxy)hydroxide. In the present study, we tested the feasibility of converting the vaccine from a liquid suspension into dry powder by thin-film freeze-drying (TFFD). With the proper amount of trehalose and/or sucrose as cryoprotectant, TFFD can be applied to transform the Norovirus vaccine candidate into dry powders without causing antigen loss or particle aggregation, while maintaining the potency of the antigens within a specified acceptable range. In an accelerated stability study, the potency of the antigens was also maintained in the specified acceptable range after the dry powders were stored for eight weeks at 40°C, 75% relative humidity. The dry powder Norovirus vaccine offers the potential to eliminate the cold chain requirement for transport and/or storage of the vaccine.

## 1. Introduction

The World Health Organization (WHO) recommends that the majority of the human vaccines should be kept in cold chain storage of 2 to 8°C during transport and storage (1). Many vaccines such as Prevnar-13 and Gardasil-9 contain an insoluble aluminum salt as an adjuvant. Unfortunately, vaccines adjuvanted with aluminum salt must not be exposed to freezing temperatures during transport or storage, because the unintentional slow freezing causes irreversible aggregation of the insoluble aluminum salt microparticles, leading to a loss of vaccine potency and efficacy (2-5). In addition, the protein antigens in the liquid vaccines are too fragile to be stable when exposed for an extended period of time to room temperatures or above. The cold chain requirement significantly hinders the distribution of vaccines globally because it brings costly waste to the vaccine supply (6, 7), and inadvertent exposure of the vaccines to freezing or heat could compromise their potency, potentially leading to inadvertent administration of vaccines with suboptimal potency or efficacy (4, 8-11).

Thin-film freezing (TFF) is a rapid freezing technique, with a freezing rate of 100-1000 K/s (12). Thin-film freeze-drying (TFFD) has proven a viable method to transform vaccines containing an insoluble aluminum salt from a liquid suspension to dry powder (13-15). Previously, we have demonstrated the feasibility of applying TFFD to several different vaccines, including model vaccines as well as commercially available veterinary and human vaccines that contain aluminum (oxy)hydroxide, aluminum (hydroxy)phosphate, potassium aluminum sulfate, or amorphous aluminum hydroxyphosphate sulfate (e.g. tetanus toxoid vaccine, Engerix-B human hepatitis B vaccine, and Gardasil human papillomavirus vaccine) (13). It was showed that subjecting these vaccines to TFFD did not cause particle aggregation after reconstitution (13). Moreover, in a mouse model, it was shown that thin-film freeze-dried vaccine was as immunogenic as the vaccine before it was subjected to TFFD (13). Importantly, the resultant dry power vaccine was not sensitive to multiple cycles of freezing (in a -20°C freezer)-and-thawing (13), or to temperatures as high as 40°C for an extended period of time (15). For example, the stability of ovalbumin (OVA) adsorbed onto aluminum (oxy)hydroxide, a model vaccine, after subjected to TFFD was evaluated. The immunogenicity of the dry powder vaccine did not significantly change after it was stored for three months at room temperature (24-25°C) or 40°C, whereas the same vaccine in a liquid suspension almost completely lost its immunogenicity after three months of storage at room temperature (15).

Norovirus has been indicated as the one of the most common causes of acute gastroenteritis in subjects of any age (16). It was reported that Norovirus causes nearly 700 million cases of illness with significant morbidity, which leads to a worldwide social burden (i.e. a total of $4.2 billion estimated in direct health system costs and $60.3 billion in societal costs per year) (16). More than 200,000 deaths per year are estimated to result from Norovirus illness, primarily in developing countries (17), where a vaccine that does not require cold chain for storage and transport would be most beneficial and essential. In 2016, the WHO stated that the development of a Norovirus vaccine should be an absolute priority. A liquid Norovirus vaccine candidate was developed for intramuscular injection to cover the two genogroups that cause the majority of illness in humans (18, 19). Specifically, it is a bivalent virus like particle (VLP) vaccine and consists of antigens from two Norovirus strains, GI.1 Norwalk and GII.4 Consensus, adsorbed on aluminum (oxy)hydroxide in suspension. Data from a phase 1/2 study showed that this vaccine candidate is generally well-tolerated and has a clinically relevant impact on the symptoms and severity of Norovirus illness after challenge (18, 19). In the present study, we tested the feasibility of applying TFFD to transform the liquid Norovirus vaccine candidate into dry powders and evaluated the potency of the antigens in the vaccine dry powders in an accelerated stability study.

## 2. Materials and Methods

### 2.1. Materials

The Norovirus vaccine is a bivalent VLP vaccine and consists of Norovirus strain GI.1 Norwalk VLP and strain GII.4 Consensus VLP adsorbed on aluminum (oxy)hydroxide. The final drug product formulation contains 50 μg of GI.1 Norwalk VLP, 150 μg GII.4 Consensus VLP, and 500 μg aluminum (oxy)hydroxide per 0.5 mL dose. The bulk drug product also contains 20 mM of L-histidine, 150 mM of NaCl, and 6.11 mg/mL of sucrose at pH 6.6-7.0. Sucrose (low in endotoxins, suitable for use as excipient EMPROVE® exp Ph Eur, BP, JP, NF) and methanol anhydrate were from Sigma-Aldrich (St. Louis, MO). Trehalose dihydrate (USP grade) was from Pfanstiehl (Waukegan, IL). Aluminum pouches were from IMPAK (Los Angeles, CA). Desiccant was from W.A. Hammond Drierite (Xenia, OH). Silanized 20 mL glass vials were from Schott (Mainz, Germany).

### 2.2. Preparation of vaccine formulations for thin-film freeze-drying

The Norovirus vaccine candidate was mixed with a sucrose or trehalose stock solution in water to reach a final sucrose concentration of 0.55-5.55% (w/v), or a final trehalose concentration of 0-5% (w/v). It is noted the vaccine formulations containing various concentration of trehalose also contained sucrose at 0.55% (w/v), as the liquid Norovirus vaccine candidate contains 6.11 mg/mL of sucrose already.

### 2.3. Thin-film freezing, sublimation, and packaging

The Norovirus vaccine in suspension containing sucrose (0.55% - 5.55%, w/v) or trehalose (0-5%, w/v) and sucrose at 0.55% (w/v) was subjected to TFF (12, 13, 20). Bulk TFF process was done as previously described (13, 21-23). Briefly, vaccine suspension was applied dropwise onto a rotating cryogenically cooled stainless steel surface where it is frozen into thin-films. The rotating speed at which the cryogenic steel surface was controlled at 5–7 rpm to avoid the overlap of films. The frozen thin-films were removed by a steel blade and collected in liquid nitrogen in a salinized glass. The glass vial was capped with rubber stopper with half open and then transferred into a VirTis Advantage bench top tray lyophilizer with stopper re-cap function (The VirTis Company, Inc. Gardiner, NY).

For single-vial TFF, a salinized glass vial was immersed into liquid nitrogen for 10 min to create a cryogenically cooled surface in the inner bottom of the vial, without the liquid nitrogen entering the vial. A syringe with a 18G1 needle was used to add 250 μL of the Norovirus vaccine candidate in suspension dropwise to the bottom of the vial so that the droplets, upon impact of the surface, rapidly froze into thin-films. The vial was capped with a rubber stopper with half open and then transferred to the lyophilizer as mentioned above for lyophilization.

Sublimation was performed over 60 h at pressures less than 200 mTorr, while the shelf temperature was gradually ramped from -40°C to 25°C. The primary drying time and secondary dry time were both 20 h. After lyophilization, vacuum was released, and nitrogen gas was filled inside the lyophilizer. The glass vial was capped tightly with rubber stopper using the automatic stopper-recap function in the lyophilizer and sealed with aluminum cap. The vial was then placed individually into an aluminum pouch with desiccant inside. The pouch was sealed, shipped from The University of Texas at Austin (UT Austin) to Takeda Vaccine in 2-8°C and then stored in a 4°C refrigerator before further use.

### 2.4. Characterization of the thin-film freeze-dried Norovirus vaccine powders

The moisture content in the dried powder was determined using a Karl Fisher Titrator Aquapal III from CSC Scientific Company (Fairfax, VA). To determine the particle size and size distribution, vials with thin-film freeze-dried Norovirus vaccine powders were randomly selected, and the powder was reconstituted with water (1 mL for bulk TFFD, and 250 μL for single-vial TFFD). The particle size and size distribution in the reconstituted Norovirus vaccine, after 2-fold dilution to reach an obscuration effect of ∼10%, were determined using a Sympatec HELOS laser diffraction instrument equipped with a R3 lens (Sympatec GmbH, Germany). The pH value of the reconstituted Norovirus vaccine was measured using a S220 SevenCompact pH meter (Mettler Toledo, OH). The concentrations of the GI.1 Norwalk VLP and GII.4 Consensus VLP antigens (total VLP and adsorbed VLP) in the reconstituted vaccines were determined using an Agilent 1260 RP-HPLC equipped with an Agilent PoroShell 300SB-C8 2.1 × 75 mm, 5 µm (Agilent Technologies, Santa Clara, CA). The following HPLC conditions were used: mobile phase A 0.1% trifluoroacetic acid (TFA) in water, mobile phase B 0.06% TFA in acetonitrile, flow rate 1 mL/min, detector wave lengths 220/280 nm. The antigen adsorption efficient value was calculated by dividing the amount of adsorbed antigen with the total antigen amount. The potency of the antigens before and after the vaccine was subjected to TFFD and reconstitution was determined using an in vitro relative potency (IVRP) assay. Briefly, 96-well plates were coated with either GII.4 Consensus or GI.1 Norwalk reference VLPs. Reference Vaccine, Test Vaccine, and assay control sample (ACS) were serially diluted in a dilution plate in the presence of goat serum and incubated with a fixed concentration of specific monoclonal antibodies (mAbs) to either GII.4 Consensus or GI.1 Norwalk VLPs. Dilution plate was transferred to VLP-coated plate. Unbound mAb remaining after incubation was allowed to bind to the VLPs on the bottom of the plate. Secondary antibodies were used to detect mAb and dilution curves are fitted using 4-Parameter Logistic (4-PL) fit. The OD value was read using a SpectraMax i3x plate reader from Molecular Devices LLC (San Jose, CA) capable of reading absorbance at 405 nm; The plate washer was a Molecular Devices’ AquaMax 4000; The incubator was from Thermo Scientific with setting of 37°C, ≥ 85% relative humidity, 5% CO_2_. Relative potency (RP) was determined for each VLP using the following equation:

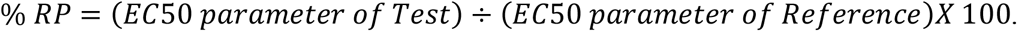

RP was normalized to antigen concentration determined using HLPC. Based on the variability of the assay, the specified acceptable potency range was 50%-200%.

### 2.5. Accelerated stability study

Randomly selected vials with thin-film freeze-dried Norovirus vaccine powders were stored at 40°C, 75% relative humidity (RH), and the potency of the antigens upon reconstitution after 4 weeks and 8 weeks of storage was determined using the IVRP assay as mentioned above.

## 3. Results and Discussion

### 3.1. Effect of the concentration of trehalose or sucrose on the particle size and size distribution of Norovirus vaccine after it was subjected to TFFD and reconstitution

A wide range of excipients such as sugars, sugar alcohols, amino acids, polymers, and/or proteins are typically used when freeze-drying vaccines containing an aluminum salt (24-27). Nonreducing sugars, such as trehalose and sucrose, with relatively high glass transition temperatures (i.e. ∼105-120°C and 60-70°C, for pure trehalose and pure sucrose, respectively) are typically used as the primary excipients (28, 29). Upon drying, these sugars form a glass as opposed to crystals to maximize their stabilization effect (30). Trehalose and sucrose are effective stabilizers. They reportedly act by hydrogen bonding to the dried antigens and/or the aluminum salt microparticles, as a water substitute (31). It is proposed that the amorphous character of the glass enables the intimate contact required for the formation of hydrogen bonds to occur between the sugars and proteins, and likely between the sugars and the antigen-adsorbed aluminum salt microparticles (32, 33). Alternative hypotheses, including vitrification (34) and particle isolation (35), have also been proposed to explain how glass-forming stabilizers preserve antigens as well as the aluminum salt microparticles. Data from our previous studies showed that trehalose at a concentration at low as 2% was sufficient to protect several vaccines adjuvanted with aluminum salts when they were subjected to TFFD (13). In the present study, trehalose and sucrose were selected as primary excipients when thin-film freeze-drying the Norovirus vaccine. It was hypothesized that they would stabilize the antigens, GI.1 Norwalk VLP and GII.4 Consensus VLP, and help preserve their potency and prevent the aggregation of the antigen-adsorbed aluminum (oxy)hydroxide microparticles in the vaccine during the thin-film freeze-drying process.

When measuring particles size and size distribution in the Norovirus vaccine, to confirm consistency and reproducibility in the readings, the effect of the obscuration value on the particle size data was studied first by determining the particle size and size distribution after the original Norovirus vaccine was diluted in water. It was concluded that the vaccine needed to be diluted two-fold to reach an optimum obscuration level of no more than 10-15%, based on the recommended range for the material size measurement (36). Shown in Figure 1A is the effect of concentration of trehalose on the particle size and size distribution after the Norovirus vaccine was subjected to bulk TFFD and reconstitution, while Figure 1B shows the effect of the concentration of sucrose on the particle size and size distribution after the Norovirus vaccine was subjected to bulk TFFD and reconstitution. Trehalose at 4% (w/v, with 0.55% (w/v) of sucrose) or sucrose alone at 4.55-5.55 % (w/v) were needed to help maintain the particle size and size distribution in the Norovirus vaccine after it was subjected to TFFD, because these formulations showed no or minimum change in particle size and size distribution.

**Figure 1.**
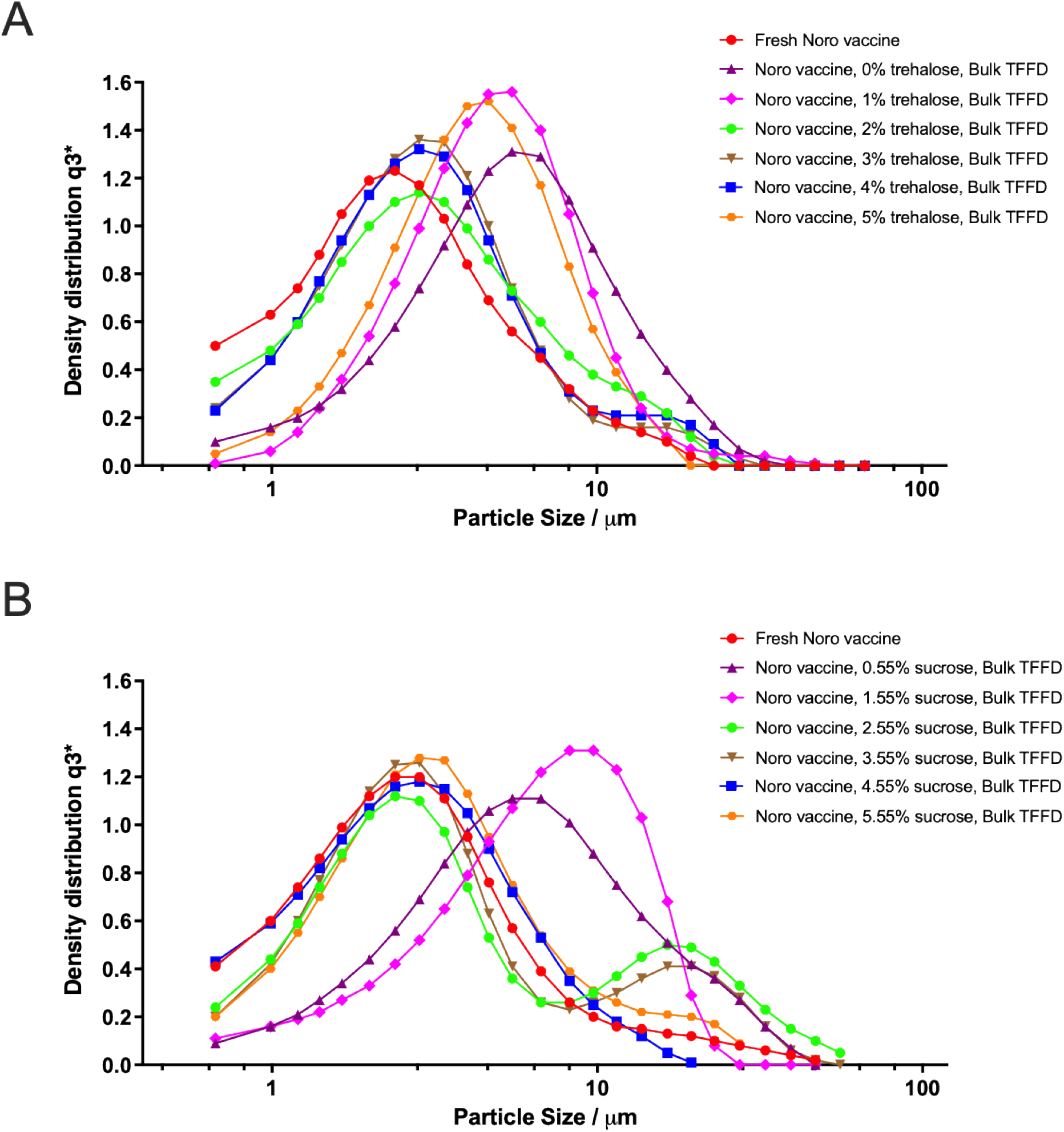
Representative particle size distribution curves of Norovirus vaccine reconstituted from powders that were prepared by thin-film freeze-drying using various concentrations of trehalose (i.e., 0–5%, w/v, with 0.55% of sucrose) (A) or sucrose alone (i.e., 0.55–5.55%, w/v) (B). The measurements were repeated three times with similar results.

In liquid vaccines that contain an insoluble aluminum salt as an adjuvant, water molecules form a hydration shell surrounding the antigens and likely the antigen-adsorbed aluminum salt microparticles, helping to preserve the antigen in its native form and the aluminum salts in a colloidal suspension. During a slow freezing process (e.g., shelf freezing in a conventional freeze-drying process, or unintentional freezing during transport or storage), the few large ice crystals that form disrupt the hydration shell by depleting water molecules, resulting in antigen unfolding and adjuvant aggregation. The extent of this irreversible aggregation is dependent on parameters such as the freezing rate (2). The ability of the TFFD technology to help prevent aggregation of the antigens and the antigen-adsorbed aluminum (oxy)hydroxide microparticles during the thin-film freezing and subsequent sublimation processes was likely due to the ultra-rapid freezing (i.e., ∼100-1000 K/s, or a time scale of 70-1000 ms) (12). The high freezing rate helps to rapidly create numerous, small nucleated ice crystals, leaving very thin channels between the frozen ice crystals. The thin channels limit the chance of collisions between antigen proteins and/or antigen-adsorbed aluminum salt microparticles and thus reduce their aggregation (12). In addition, the trehalose or sucrose in the vaccine colloidal suspension may not only replace water molecules to bind to the protein antigens or the antigen-adsorbed aluminum salt microparticles, but also help to increase the viscosity in the thin channels, further reducing the mobility of the antigen proteins or the antigen-adsorbed aluminum salt microparticles and thus reduce the chance for them to collide and aggregate (12).

### 3.2. Effect of bulk TFFD vs. single-vial TFFD on the particle size and size distribution of Norovirus vaccine

To understand the effect of the bulk TFF process compared to the single-vial TFF process on the particle size and size distribution of Norovirus vaccine after TFFD, Norovirus vaccine suspended in 4% (w/v) of trehalose (with 0.55%, w/v, of sucrose) or in 5.55% (w/v) sucrose alone was subjected to single-vial TFFD or bulk TFFD. Shown in Figure 2 and Table 1 are representative particle size and size distribution after the dry powders were reconstituted. Overall, it appeared that subjecting the Norovirus vaccine to either single-vial TFFD or bulk TFFD did not cause significant particle aggregation, demonstrating the versatility of the TFFD technology as the vaccine may be directly thin-film frozen in individual vials to form frozen thin-films, and the water molecules are then removed in the frozen thin-films by sublimation. Otherwise, the vaccine may also be bulk frozen into frozen thin-films. After the water molecules in the frozen thin-films are removed by sublimation, the resultant bulk dry powder can then be filled into containers such as glass vials.

**Table 1.** Particle size and size distribution of Norovirus vaccine after it was subjected to bulk or single thin-film freeze-drying in 4% (w/v) trehalose (with 0.55% (w/v) of sucrose) or 5.55% (w/v) sucrose alone and then reconstitution.

| Formulations | X<sub>10</sub> (μm) | X<sub>50</sub> (μm) | X<sub>90</sub> (μm) |
| --- | --- | --- | --- |
| Norovirus vaccine | 1.02 ± 0.19 | 2.71 ± 0.33 | 7.67 ± 0.17 |
| Norovirus vaccine, 5.55% sucrose, bulk | 1.13 ± 0.21 | 2.62 ± 0.42 | 8.65 ± 2.74 |
| Norovirus vaccine, 5.55% sucrose, single vial | 0.99 ± 0.16 | 2.85 ± 0.45 | 6.76 ± 0.13* |
| Norovirus vaccine, 4% trehalose/0.55% sucrose, bulk | 1.62 ± 0.26 | 3.82 ± 0.57 | 8.43 ± 0.67 |
| Norovirus vaccine, 4% trehalose/0.55% sucrose, single via | 1.52 ± 0.27 | 3.84 ± 0.45 | 8.86 ± 2.72 |

X<sub>10</sub>, X<sub>50</sub>, X<sub>90</sub> denote particle dimensions corresponding to 10%, 50% and 90% of the cumulative undersize distribution. Data are mean ± SD (n = 2 or 3. \**p* < 0.05, vs. fresh, T-test, two tails).

**Figure 2.**
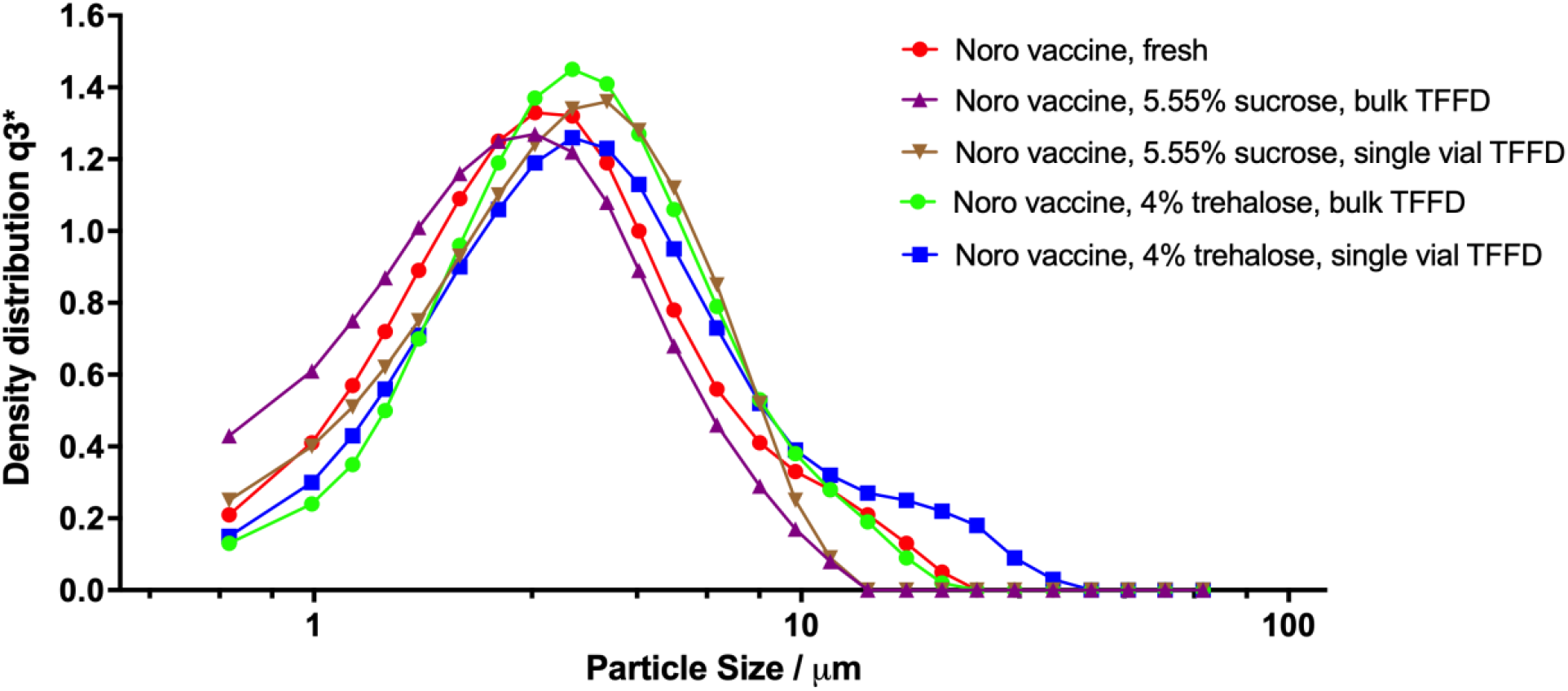
Representative particle size distribution curves of bulk and single vial thin-film freeze-dried Norovirus vaccine powders after reconstitution. The vaccine formulations contained 5.55% (w/v) of sucrose alone or 4% (w/v) of trehalose with 0.55% (w/v) of sucrose. The measurement was repeated with two or three vials with similar results.

### 3.3. Characterization of thin-film freeze-dried Norovirus vaccine

It is a mandatory requirement to specify the water content of all dry powder vaccines prepared for clinical use (37). Among the currently licensed and commercially available vaccines, notably many live-attenuated vaccines are particularly unstable unless stored as a dry product with a low residual moisture content (typically less than ∼3%) (38). Dry powder vaccines with high moisture content can result in poor stability since free water molecules induce conformational changes in protein antigens, similar to those experienced in a liquid solution (39). In vaccines containing aluminum salts as the adjuvant, large amount of residual free water molecules in the powder will also allow the aluminum salt microparticles to move and collide, ultimately leading to aggregation. Once the dry powder vaccine has been reconstituted, potency often decreases rapidly, especially if the product is not kept cold (1). The residual moisture content in the thin-film freeze-dried Norovirus vaccine powders were 1.5-1.8% (Table 2), well below the suggested 3% limit (38).

**Table 2.**
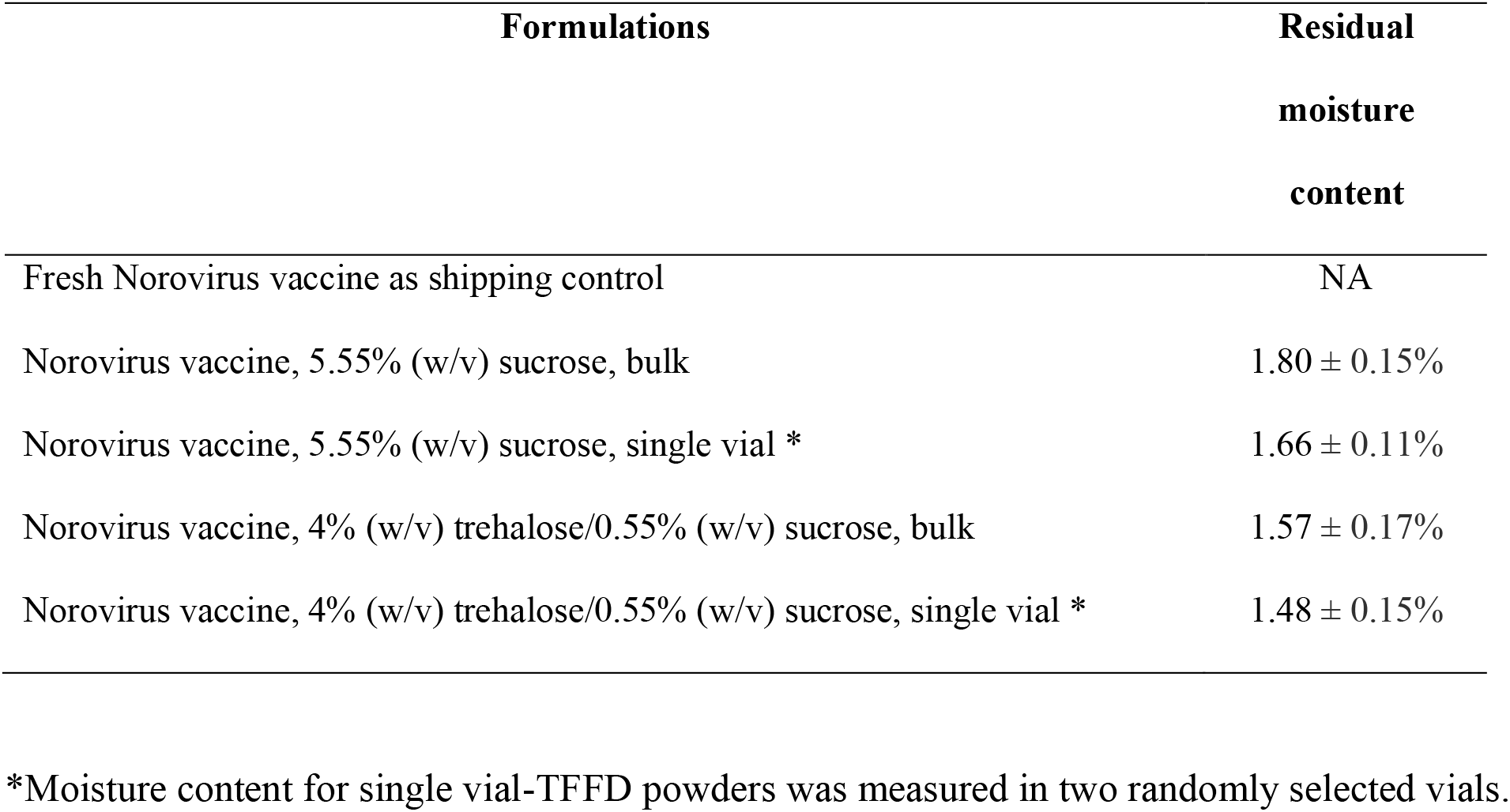
Residual moisture content in thin-film freeze-dried Norovirus vaccine powders. Data are mean ± S.D.

Upon reconstituting the Norovirus vaccine dry powders prepared with 4% (w/v) of trehalose (with 0.55%, w/v, of sucrose) or 5.55% (w/v) of sucrose alone by either single-vial TFFD or bulk TFFD, we also measured the pH, antigen concentration, and antigen potency in the reconstituted Norovirus vaccine suspension. The pH values of the vaccine remained 6.4-6.5. The recovery rate of the GII.4 Consensus VLP antigen was in the range of 82.2-118.1%, 93.8-130.4% for the GI.1 Norwalk VLP antigen, as determined by RP-HPLC. The percentage of antigen adsorbed for the GII.4 Consensus VLP antigen was 97.7-103.1%, 98.4%-103.1% for the GI.1 Norwalk VLP antigen. The relative potency as measured by the IVRP assay for the Consensus and Norwalk VLP antigens are shown in Figure 3. The relative potency of the Norwalk VLP antigen in all four formulations met the specified acceptable range of 50%-200% (Fig. 3A, i.e. 101.2%-171.8%). For the Consensus VLP antigen, all formulations, except the one containing 4% (w/v) trehalose and 0.55% (w/v) of sucrose and prepared using the bulk TFFD method, also met the specified acceptable range (Fig. 3B). The potency of both antigens appeared to be better maintained in Norovirus vaccine powders prepared with sucrose at 5.55% (w/v), especially when thin-film freeze-dried using the single vial method (Fig. 3), likely because during single vial TFFD, all the frozen thin-films were recovered, while in the bulk TFFD, a small percent of the frozen thin-films may not be collected when they were transferred from the cryogenically cooled stainless steel surface into a glass vial, which could be minimized in the future by adjusting the filling of the frozen thin films or the resultant dry powders. Overall, it is concluded that the Norovirus vaccine can be transformed into a dry powder using TFFD while preserving particle size and size distribution of the antigen-adsorbed aluminum (oxy)hydroxide microparticles as well as the chemical integrity and potency of the antigens in the specified acceptable range.

**Figure 3.**
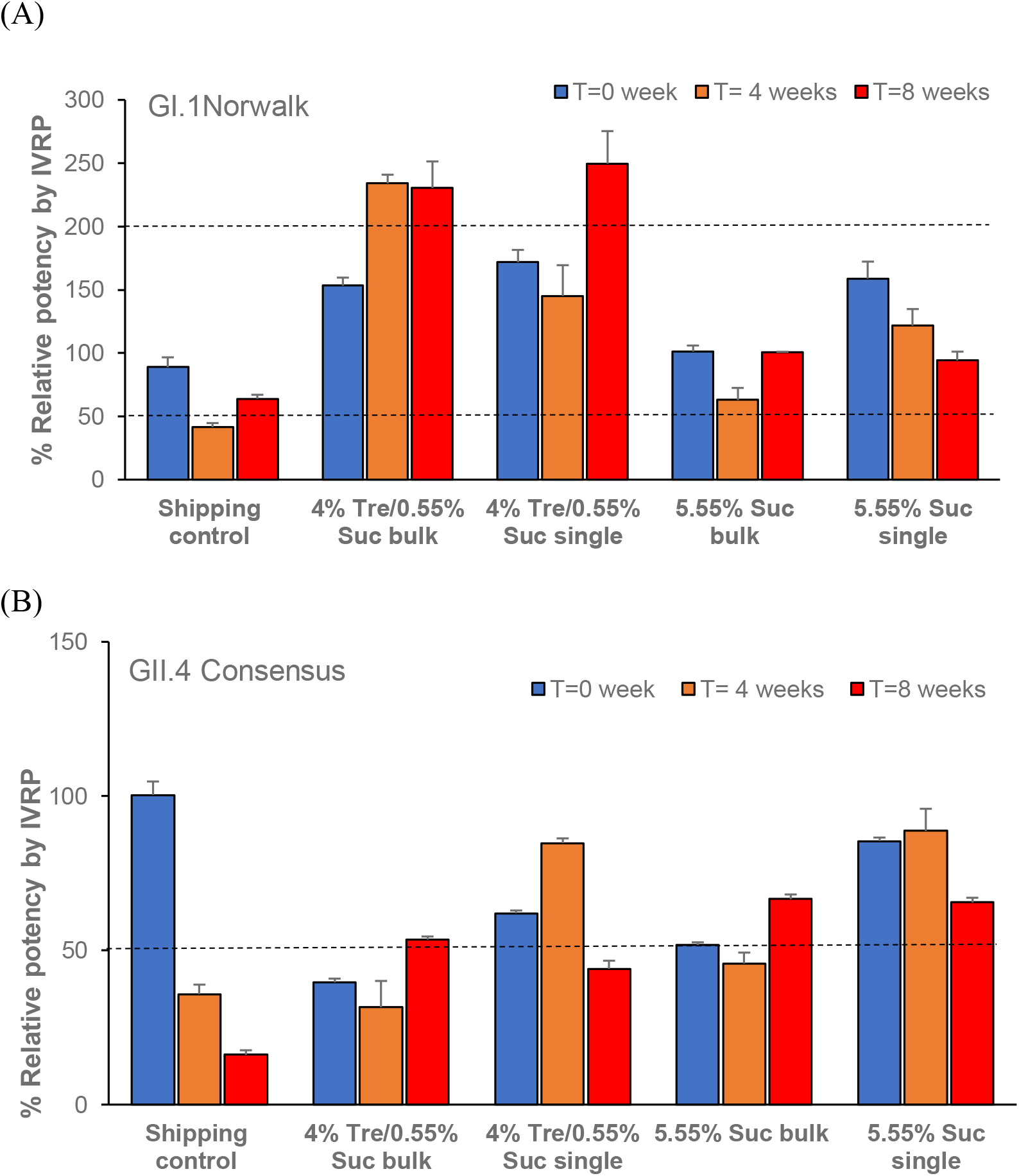
Potency of antigens, GI.1 Norwalk VLP (A) and GII.4 Consensus VLP (B), in thin-film freeze-dried Norovirus vaccine powders after 0, 4, or 8 weeks of storage at 40°C, 75% relative humidity. Data are mean ± SD (n = 3). For the IVRP assay, based on the variability of the assay, relative potency in the range of 50%-200% (see dashed horizontal lines) is acceptable.

### 3.4. Stability of the Norovirus vaccine after eight weeks of storage as thin-film freeze-dried powders at 40°C, 75% relative humidity

The importance of vaccine stability has been acknowledged and emphasized by the WHO. In vaccine evaluation, therefore, many different stability indicating assays are used (40, 41). The antigen potency is one of most important characteristics in studying the thermostability of vaccines (1). The WHO guidelines in 2006 stressed the need for stability data to support clinical trial approval for vaccines (42). In an accelerated stability study, we stored the Norovirus vaccine dry powder at 40°C, 75% RH, for 8 weeks and measured the antigen content and integrity, antigen adsorption efficiency to the aluminum salt, and antigen potency in week 4 and week 8. As shown in Figure 3A, the relative potency of GI.1 Norwalk VLP antigen in all four dry powder formulations remained within the specified acceptable range of 50-200% after 8 weeks of storage. For GII.4 Consensus VLP antigen, the relative potency for the dry powder formulations containing 5.55% (w/v) sucrose prepared by either bulk TFFD or single vial TFFD also remained within the acceptable range after 8 weeks of storage. However, for the formulations containing trehalose (4%, w/v) and sucrose 0.55% (w/v), the relative potency reduced to below 50% (Fig. 3). Moreover, the percentage of antigen adsorbed for both antigens remained close to 100% after all dry powders were subjected to 8 weeks of storage at 40°C, 75% RH (i.e. 102.1%-111.9% for Norwalk VLP antigen, and 100.7%-105.2% for Consensus VLP antigen). As for the control, i.e. that liquid vaccine that was shipped at 2-8°C from Takeda Vaccine to UT Austin and from UT Austin to Takeda Vaccine together with the dry powders (in separate vials) and then subject to the same 40°C, 75% RH storage condition, the relative potency values of both Norwalk VLP and Consensus VLP antigens were reduced to below 50% after the 8 weeks of storage at 40°C, 75% RH. Taken together, the potency of the antigens in the Norovirus vaccine after subject to TFFD with the proper excipients, e.g. sucrose at 5.55% (w/v), was maintained within the specified acceptable range, and the potency remained within the acceptable range even after the dry powders prepared with 5.55% sucrose (w/v) in an accelerated stability study, indicating the potential of storing the thin-film freeze-dried Norovirus vaccine powder without the 2-8°C cold chain or at least in a controlled temperature chain (CTC). It is likely that the sucrose and/or trehalose in the Norovirus vaccine powders formed hydrogen bonds to the dried antigens or the antigen-adsorbed aluminum salt microparticles and acted as a water substitute to stabilize the vaccine (31). The sucrose and trehalose also likely acted as glass-forming stabilizers to vitrify and isolate the protein antigens and/or the antigen-adsorbed aluminum salt microparticles in the dry powder (34, 35), minimizing the mobility of the antigens and particles and thus preventing their collision, even in a temperature as high as 40°C. The low moisture content in the thin-film freeze-dried Norovirus vaccine powders as well as the package strategy (i.e., filling vials with nitrogen gas, capping vials with rubber stoppers and aluminum crimp, and sealing of the vial individually in an aluminum pouch containing desiccant) likely have also contributed to the stability of the Norovirus vaccine dry powders during transport and storage.

## Conclusion

Thin-film freeze-drying was applied to successfully transform a bivalent Norovirus vaccine candidate adjuvanted with an insoluble aluminum salt from a liquid suspension to dry powders without causing particle aggregation or antigen loss, while maintaining the potency of the antigens within our acceptable range. Importantly, relative potency of Norovirus vaccine antigens, both Norwalk VLP and Consensus VLP, in the thin-film freeze-dried powders prepare with 5.55% (w/v) of sucrose remained within the acceptable range in an accelerated stability study.

## Acknowledgements

This work was supported by Takeda Vaccine via a Technology Validation Agreement between UT Austin and Takeda Vaccine.

